# The Collaborative Cross Graphical Genome

**DOI:** 10.1101/858142

**Authors:** Hang Su, Ziwei Chen, Jaytheert Rao, Maya Najarian, John Shorter, Fernando Pardo Manuel de Villena, Leonard McMillan

## Abstract

The mouse reference is one of the most widely used and accurately assembled mammalian genomes, and is the foundation for a wide range of bioinformatics and genetics tools. However, it represents the genomic organization of a single inbred mouse strain. Recently, inexpensive and fast genome sequencing has enabled the assembly of other common mouse strains at a quality approaching that of the reference. However, using these alternative assemblies in standard genomics analysis pipelines presents significant challenges. It has been suggested that a pangenome reference assembly, which incorporates multiple genomes into a single representation, are the path forward, but there are few standards for, or instances of practical pangenome representations suitable for large eukaryotic genomes. We present a pragmatic graph-based pangenome representation as a genomic resource for the widely-used recombinant-inbred mouse genetic reference population known as the Collaborative Cross (CC) and its eight founder genomes. Our pangenome representation leverages existing standards for genomic sequence representations with backward-compatible extensions to describe graph topology and genome-specific annotations along paths. It packs 83 mouse genomes (8 founders + 75 CC strains) into a single graph representation that captures important notions relating genomes such as identity-by-descent and highly variable genomic regions. The introduction of special anchor nodes with sequence content provides a valid coordinate framework that divides large eukaryotic genomes into homologous segments and addresses most of the graph-based position reference issues. Parallel edges between anchors place variants within a context that facilitates orthogonal genome comparison and visualization. Furthermore, our graph structure allows annotations to be placed in multiple genomic contexts and simplifies their maintenance as the assembly improves. The CC reference pangenome provides an open framework for new tool chain development and analysis.

## Introduction

Reference genome assemblies are the foundation of most bioinformatics pipelines. They provide coordinate frames and serve as the substrate for discovery and annotation. Genome assemblies enable downstream computational analyses including searching, mapping, and comparison. Reference genomes are also a critical resource for genotyping and assay design. Most existing reference assemblies model either a single representative, a population consensus, a minimal “conserved” genome, or a “maximal” genome that attempts to capture all known variants (Computational Pan-Genomics 2018). While high quality reference assemblies are available, it is well recognized that using a single reference assembly to represent a population comes with limitations since reference genome have gaps, mis-assigned regions, and do not capture all of the genetic diversity within a species (Church et al. 2011; Church et al. 2015; Computational Pan-Genomics 2018). The reliance on a single reference assembly leads to issues such as poor characterization or loss of genomic regions with complex and variable structural diversity (Dilthey et al. 2015) or allele-specific function (Stevenson et al. 2013). Moreover, in the face of the growing numbers of available genome assemblies enabled by modern high-throughput sequencing technologies development, bioinformaticians are confronted with a choice of either selecting one from multiple assembly choices or performing multiple analyses and somehow merging their results.

Pangenomes have been suggested as the next logical step for representing reference genome assemblies (Church et al. 2015; Paten et al. 2017; Computational Pan-Genomics 2018). A pangenome is defined as a collection of genomic sequences that are analyzed and represented together as a reference to achieve *completeness, stability, comprehensiveness, and efficiency* (Computational Pan-Genomics 2018). Most reference pangenomes are represented by graphs in which alternate paths either model genomic variants or differences in genome organization. Here, we will distinguish a *reference* pangenome as being population specific (Li et al. 2010), in contrast to either an inter-species *phylogenetic* pangenome that represents an evolutionary history of presently diverged clades (Tettelin et al. 2005; Tettelin et al. 2008), or a *mutagenic* pangenome that represents somatic DNA variations within a single organism such as those proposed for cancer study (Zheng et al. 2016). Moreover, we refer to traditional *single sequence* reference assemblies as *linear* reference genomes. Reference pangenomes are already available for viral and prokaryotic strains and populations (Tettelin et al. 2005; Hogg et al. 2007; Lefebure and Stanhope 2007; Jacobsen et al. 2011; Zhou et al. 2018), but are less common for eukaryotes (Li et al. 2010; Hu et al. 2017; Peter et al. 2018; Tian et al. 2018; McCarthy and Fitzpatrick 2019). Common pangenome representations includes multiple sequence alignments, kmer-based, or graphs. Common objectives of pangenome representations include minimizing redundancy (Marcus et al. 2014; Baier et al. 2015), estimating genome size (Lapierre and Gogarten 2009), supporting the addition of new genomes (Paten et al. 2017), and establishing coordinates for annotations (Herbig et al. 2012; Rand et al. 2017; Gartner et al. 2018). A comprehensive reference pangenome that is compatible with the downstream tool chains is also a pressing need for future genomic study.

We present a pangenome resource for a multi-parental population of widely studied mouse strains. The Collaborative Cross (CC) is a large panel of recombinant-inbred strains derived from a genetically diverse set of eight laboratory mouse strains designed specifically for complex-trait analysis (Churchill et al. 2004). The genome of each CC strain sample can be regarded as a mosaic of its eight founder genomes (Figure S1). The CC are popular genetic reference populations due to a huge expansion of phenotypic variation observed for most traits measured using them (Aylor et al. 2011; Philip et al. 2011; Bottomly et al. 2012; Kelada et al. 2012; Leist et al. 2016; Manet et al. 2019). This has motivated the creation of new CC-related genomic resources (Srivastava et al. 2017; Shorter et al. 2019) and analysis tools (Holt et al. 2013; Huang et al. 2014) to aid in sorting out the underlying phenotypic correlates to the genetic sources of variation. In the summer of 2017 the sequence data from 69 CC mouse strains were published, from a single male from each available line with 20×-30× genome coverage (Srivastava et al. 2017). In early 2019, sequence data from six additional CC strains were published (Shorter et al. 2019). Meanwhile in the fall of 2018, classical “linear” genome assemblies were released for sixteen common laboratory mouse strains, which included seven of the eight CC founders (Lilue et al. 2018). The eighth strain, C57BL6/J, was already the basis for the widely used *Mus musculus* reference genome (GRCm38) (Mouse Genome Sequencing et al. 2002; Church et al. 2009).

The release of the CC strains has precipitated a need for various “*-omics*” analysis pipelines that depend on a reference genome. These analyses are complicated by biases introduced by having the reference assembly based on one of the CC population’s founders, C57BL6/J. Moreover, there are many genomic features found in other mouse strains that are absent from the standard mouse reference (Lilue et al. 2018; Mulligan et al. 2019). Since each CC line is a unique mosaic of its eight founders, some of these limitations might be overcome by somehow performing a joint analysis (Holt et al. 2013; Huang et al. 2014; Munger et al. 2014) using the separate genome reference assemblies, like those from Lilue et al (Lilue et al. 2018). However, doing so is not a simple matter. Alternatively, one could move toward a consensus genome assembly that treats founders and CC lines more uniformly, or else develop separate assemblies for each CC strain, and then compare analogous genomic regions downstream (Holt et al. 2013). Our efforts to create a CC reference pangenome, resulting from both a dire need and a serendipitous convergence of resources, in an attempt to support both traditional and new genetics and genomics analysis tools.

Here, we present a first draft of the CCGG pangenome resource. This version was constructed by merging the standard mouse reference with *de novo* assemblies of the 7 founder genomes (Lilue et al. 2018) guided by the short-read sequence data from CC-line samples (Srivastava et al. 2017; Shorter et al. 2019). The resulting pangenome represents 83 genomes (8 founders and 75 CC strains). It is composed of 2091 full-chromosome contigs and including 431 alternative autosomes sequences describing the residual heterozygosity of the sequenced CC samples. Strain-specific linear genomes can be extracted from the CCGG pangenome resource in the form of traditional FASTA files that are compatible for traditional bioinformatics pipelines. The total sequence content in pangenome is significantly smaller than the 8 founder linear genomes from which it was derived (10,653,041,524 bases vs 20,749,378,088 bases). The CCGG also provides a framework for comparative genomic analyses, where conserved subsequences, namely *anchor nodes*, partition the genomes into homologous segments with variant haplotypes, namely *edges*, that contain all sequence diversity. Anchor nodes are unique 45-mers (in both forward and reverse complement strands) found in each of the 8 founder genome assemblies that also appear in every sequenced CC sample. Moreover, the relative-order of anchors is consistent in all 8 founder assemblies. Anchor nodes provide a coordinate framework that allows annotations to be placed in a common genomic context that is easier to maintain as the assemblies are improved. By introducing anchors in a graph-based reference genome as a sequence-based coordinate framework, we address important issues (*monotonicity*, *backward-compatibility*, and *spatiality*) necessary for relating a graph-based model to traditional reference assemblies (Rand et al. 2017; Computational Pan-Genomics 2018). The graphical structure of the CCGG with annotation provides a more comprehensive picture of the genome structure surrounding biological features, which are easy to traverse for searching, annotating, comparing and visualizing genomic features. The integration of CC sequencing data in CCGG also exposes recombination cold and hotspots and annotates potentially novel gene structures created by recombination in CC strains. The CCGG serves as a practical reference genome assembly and establishes a foundation for the development of pan-genomic tools for genomic comparison, sequence alignment, and genetic association mappings.

## Results

The sequence content of the CCGG is derived from the *de novo* genome assemblies from (Lilue et al. 2018) and the standard mouse reference assembly (Mouse Genome Sequencing et al. 2002; Church et al. 2009). We refer to these eight genome assemblies as *founder* sequences. The sequencing data of CC samples from (Srivastava et al. 2017) and (Shorter et al. 2019) were used to select anchors and label edges in the graph. Since each CC genome is a mosaic of the 8 founder genomes, we annotate paths in CCGG to provide putative genomic sequences for each CC strain based on genotype probability files inferred from previous sequencing data (Srivastava et al. 2017; Shorter et al. 2019). All the paths are constrained to pass through conserved *anchor* sequences that appear in every sequenced CC sample. Furthermore, many genomic features of the reference assembly, including genes, exons, and interspersed repeats, are annotated on the anchors and edges of the CCGG’s C57BL/6J path. We also performed pairwise alignments between each parallel edge (variant subsequences) and the reference sequence and recorded these alignments annotated as CIGAR strings on each edge. The CCGG is represented as a standard FASTA file with annotations included in the sequence headers encoded as computer-readable JSON strings. An example fragment of the CCGG is shown in Figure 1.

**Figure 1.**
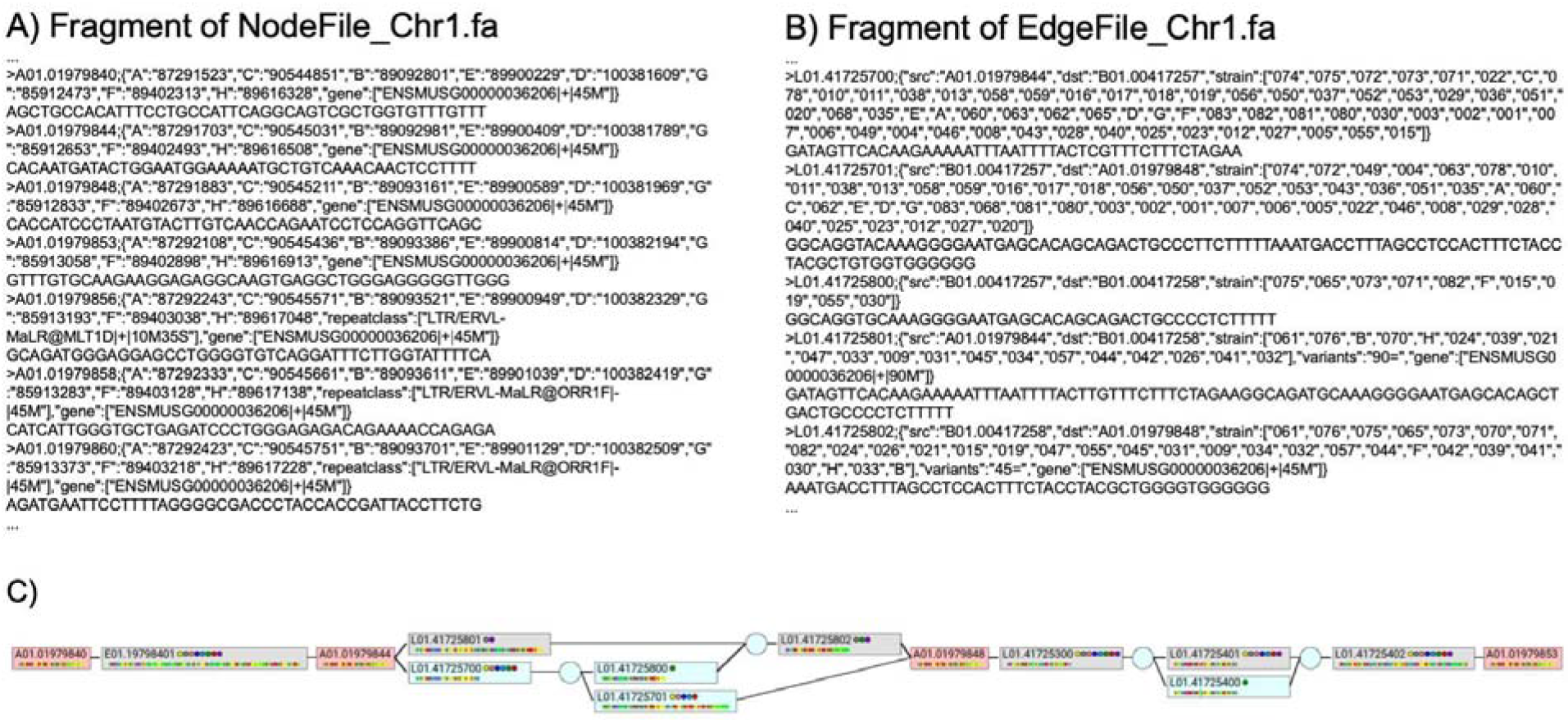
An overview of the CCGG Pan-genome file structure. The CCGG is represented as a standard FASTA file. Anchor nodes and edges are named sequences with attributes. Attributes are represented as a JSON string in the sequence’s FASTA header. A corresponding visualization of the graph fragment is shown in C. Boxes enclose both anchor nodes (pink) and edges (gray and blue), with their genomic sequences shown as a colored barcode below the node or edge name. Light gray boxes indicate edges on the GRCm38 reference path, while blue boxes indicate alternate sequences. Floating nodes, shown as blue circles, contain no sequence or annotation and only appear as source (src) or destinations (dst) in edge annotations. The strain annotation identifies the genomes with the given edge sequence where a single letter designation A-H is used for each CC founder (A/J, C57BL/6J, 129S1/SvImJ, NOD/ShiLtJ, NZO/HlLtJ, CAST/EiJ, PWK/PhJ, and WSB/EiJ respectively) and a three digit number indicates each sequenced CC sample. Anchor nodes include the linear coordinates of the anchor’s sequence in each of the founder genomes. A variant attribute is included for edges parallel to a reference sequence. The variants are indicated by a CIGAR string describing an alignment of the edge to the reference. Gene, exon, and repeat-masker annotations may appear on both anchor nodes and edges, indicating the relative position between the graph entities and these biological features. The orientations of these genomic features and their extents (described as a CIGAR string) are also provided.

### General Properties

The CCGG pangenome is a directed graph composed of nodes and edges. There are four types of nodes: *anchor nodes* which are unique and consistently ordered 45-mers common to all founder genome assemblies and are also conserved in every sequenced CC sample (i.e. their presence is supported by multiple reads), *source* and *sink* nodes represent the start and end of every contig, and *floating* nodes denote either the separation or merge of sequences between subsets of strains between adjacent anchors. Edges represent the sequences between graph nodes. Only anchor nodes and edges contain genomic sequence and annotations. Source, sink, and floating nodes do not have associated sequence or annotations.

The sequences of anchors are *conserved* in every genome represented in the graph and, thus, lie on every path. Anchor nodes are also *unique* (both forward and reverse complemented) in all eight founder genome assemblies. Anchor nodes are of a uniform size and are *consistently ordered* in all founder genome assemblies. Anchors are annotated with their linear coordinates in each of the linear founder genome assemblies. Relative to anchors one can establish offsets to specific genomic features, that fulfill desirable properties of a graph coordinate system as discussed in (Rand et al. 2017; Computational Pan-Genomics 2018), including *monotonicity*, *backward-compatibility*, and *vertical/horizontal spatiality*.

Anchors normalize the various linear genome assemblies. The uniqueness of anchor sequences establishes a one-to-one mapping between multiple genomes and partition genome sequences into homologous regions. Anchors delineate specific genomic regions supporting local sequence updates. When a new genome assembly is released, only anchor nodes need to be remapped and the intervening sequences compared and potentially updated. If a new genome assembly is incorporated into the CCGG, some anchor nodes might be demoted to an edge connected to two floating nodes, since they are no longer conserved in all genomes. The incoming and outgoing edges of the demoted anchor nodes are then reconnected to the two floating nodes (Figure S2). The new edge inherits the annotations of the primary anchor node with their “*strain*” attributes updated by incorporating the new genome notation. The choice of anchor nodes is population dependent. We estimate the possible number of anchor sequences, as determined by sequentially incorporating CC samples, fits an exponentially decaying function that converges to a non-zero asymptote (details provided in supplementary data). Extrapolation of the curve suggests that the core conserved genome is approximately 27% of the total reference genome length (95% confidence interval = 27.67% - 28.16%) and should remain relatively constant for this set of founders.

Both anchor and edge sequences allow annotations. Anchors partition genomes into homologous regions while edges lie between anchor pairs and represent a haplotype. Each edge is annotated according to the one or more genomes upon which it lays. We consider two or more paths that share common source and destination nodes as *parallel*. This notion of *parallel* extends to subgraphs separated by any pair of anchor nodes as illustrated in Figure 1. Parallel sequences represents haplotypes of a homologous region and contain all variants, which serve as an alternative variant representation to the VCF files used for traditional linear genomes.

Every genomic contig is a path beginning and terminating at special nodes named *Source* and *Sink* respectively. Through traversing the graph, one can recover exactly the 8 founder linear genome assemblies from the CCGG as well as construct putative genome of each CC sample (a command-line interface is provided for extracting full genomes). Where a CC sample is homozygous there is a single path in the graph. In regions with residual heterozygousity the graph bifurcates (in the strain annotations, for example as “001a” and “001b”) and eventually rejoins. There are a total of 2091 contigs lying between “source” and “sink” nodes. These include the assemblies for each chromosome of the 8 founder strains and homozygous/heterozygous contigs of each CC strain. The subgraphs between anchor pairs can be extracted from the CCGG by traversing and constructing all paths between the bounding anchors surrounding a specified region (i.e a gene). This provides an effective method for comparing and visualizing the genome structure to identify conserved and highly variable genomic regions. Furthermore, one can traverse a CC path in the CCGG to identify recombination events.

The CCGG pangenome is serialized as one or more standard FASTA files (Figure 1). Genomic attributes and annotations for both nodes and edges are stored as a JSON string included in each sequence’s FASTA header. We provide a Python-based API for accessing, traversing, and editing the CCGG. In addition, we provide a command-line interface for extracting contigs, subpaths, and other symbolic genomic representations from the CCGG (details given in Methods). The present CCGG pangenome is neither a minimal nor unique graph. It deliberately strikes a compromise to provide functionality while making best use of existing genomic resources.

### Sequence Commonality and Strain Diversity Pattern in CCGG

CCGG contains 10,653,041,524 bp in total. It is composed of 9,212,137 anchor nodes with 414,546,165 bp and 30,252,594 edges having 10,238,495,359 bp (autosomes and chromosome X). Anchors in the graphical genome are densely distributed, though some long gaps exist. In Figure 2A, we plot the distribution of anchors over the entire genome. On average in autosomes about 10,000 anchors fall within a 2Mb bin, giving an average of 200 bp between adjacent anchor pairs. Figure 2B illustrates the distribution of the gap spacing between adjacent anchors on autosomes. We found out that 94% of the anchors are separated by fewer than 1000 bp and 50% are separated by less than 180 bp. Anchors are less dense on chromosome X as shown in the figure. We attribute this to two factors. First, there is less unique sequence on X, and, second, Anchor sequences tend to have fewer supporting reads since the sequenced samples are hemizygous (males), and, thus, effectively having half the coverage of autosomes. In comparison, the gap lengths on chromosome X is relatively longer than on autosomes.

**Figure 2.**
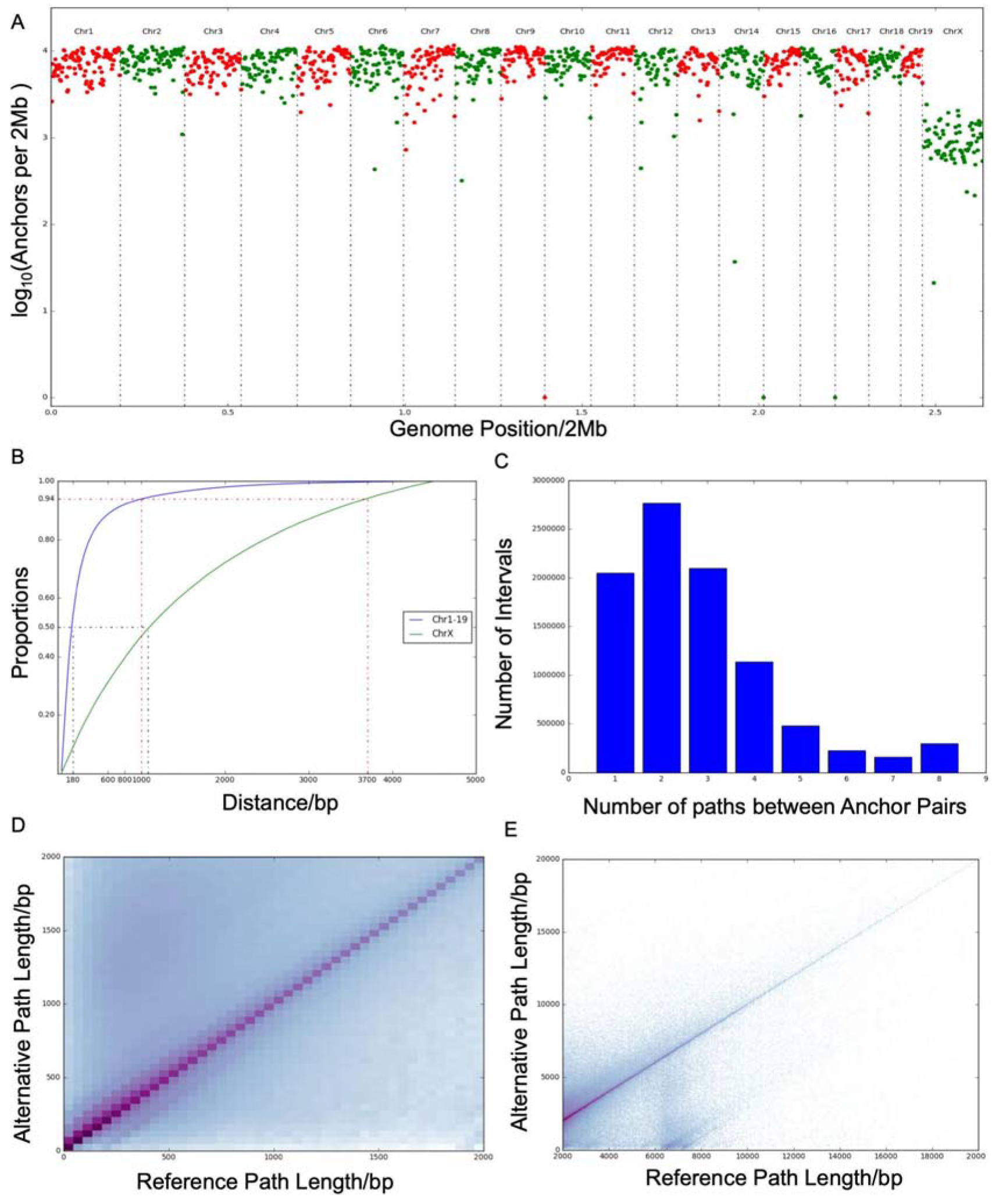
CCGG Properties. **A)** The number of anchors per 2Mb region plotted on log 10 scale. **B)** The cumulative distribution of adjacent anchor distances. The blue curve denotes the anchor separations in autosomes (chromosome 1-19), while the green curve denotes the anchor separations on chromosome X. Half of the intervals between two anchors in autosomes are within 180 bp but on chromosome X the distance is over 1000 bp. 94% of the anchors are separated by less than 1000 bp on autosomes, and 3700bp on chromosome X. **C)** The distribution of the number of separate paths between adjacent anchor nodes (chromosomes 1-19 and X) has a mode 2, which suggests many shared haplotypes exist in the CCGG. **D) Difference in Path Lengths for short anchor gaps (ref length <1000bp) as compared to the standard mouse reference.** For short gaps, the density is concentrated along the diagonal showing that reference and alternative paths are usually with similar length. **E)** For long gaps, the density is more dispersed suggesting insertions and deletions are more common in long gaps. The dense cluster below the diagonal shown in this figure for reference paths over 6000 bp suggests an excess of deletion events in alternative paths.

Anchor nodes and *collapsed edges,* defined as a single edge shared by all 8 founder genomes, represent shared sequence found in all 8 founder genomes (660,797,730 bp in total). Of 30,252,594 edges in total, 2,781,638 are collapsed edges shared by all 8 founder assemblies composed of 246,251,565 bp. The remaining edges lie on two or more parallel paths representing homologous sequences containing variants. The number of paths between adjacent anchors is shown in Figure 2C. The predominant number of paths between adjacent anchors is two (30.03%), suggesting that a large fraction of homologous regions are composed of only two haplotypes. However, there are a significant fraction of adjacent anchors with 3 (22.80%) or 4 (12.37%) parallel paths, and fewer with 5, 6, or 7, suggesting sequence variable regions exist. There is a notable increase in the number adjacent anchors with 8 parallel paths. Since long sequences tends to incorporate more variants, this increase could result from the correlation between sequence length and variants number. As shown in Figure S7, most of these 8 parallel paths appear between long gaps (80%, 229094 out of 286378 edges), but a significant fraction of these highly variant genomic regions fall between relatively short anchor gaps, suggesting high variable regions exist in the mouse genome. As shown in Figure 2D and E, most parallel edges are similar in length to the reference assembly, which suggests most common variants are either SNPs or small indels. A large variance in subpath lengths often serves as an indicator for structural variants in those regions.

For anchor intervals less than 1000 bp, we provide pairwise sequence alignments of alternative sequences and the standard mouse reference, which are annotated in a SAMtools-compatible CIGAR string on each edge called ‘*variants*’. Alignments for longer paths are generally less informative of the actual sequence variants. Thus, for long gaps where at least one path length is longer than 1000 bp, we insert floating nodes between shared sequences for further graph compression (details in Supplementary). There are 3,198,957 floating nodes in total. The length of sequences partitioned by anchors are significantly reduced by inserting floating nodes as shown in Figure S8. Floating nodes denote sharing between subsets of strains within long gaps and they create locally parallel edges. A comparison of the number of matching bases in the pairwise alignments versus the length of the parallel reference path is depicted as a 2D-histogram in Figure 3A. We also compare the number of matches and alternative path lengths in Figure 3B. As expected, the density of path-sequences is centered along the diagonal, indicating that there is significant sharing among alternative and reference paths. The lengths of reference and alternative paths is also similar as shown in Figure 2E, indicating large insertions or deletions are rare. Overall, these results suggest that, for short gaps less than 1000 bp, the alternative paths are dominated by SNPs or small indels.

**Figure 3.**
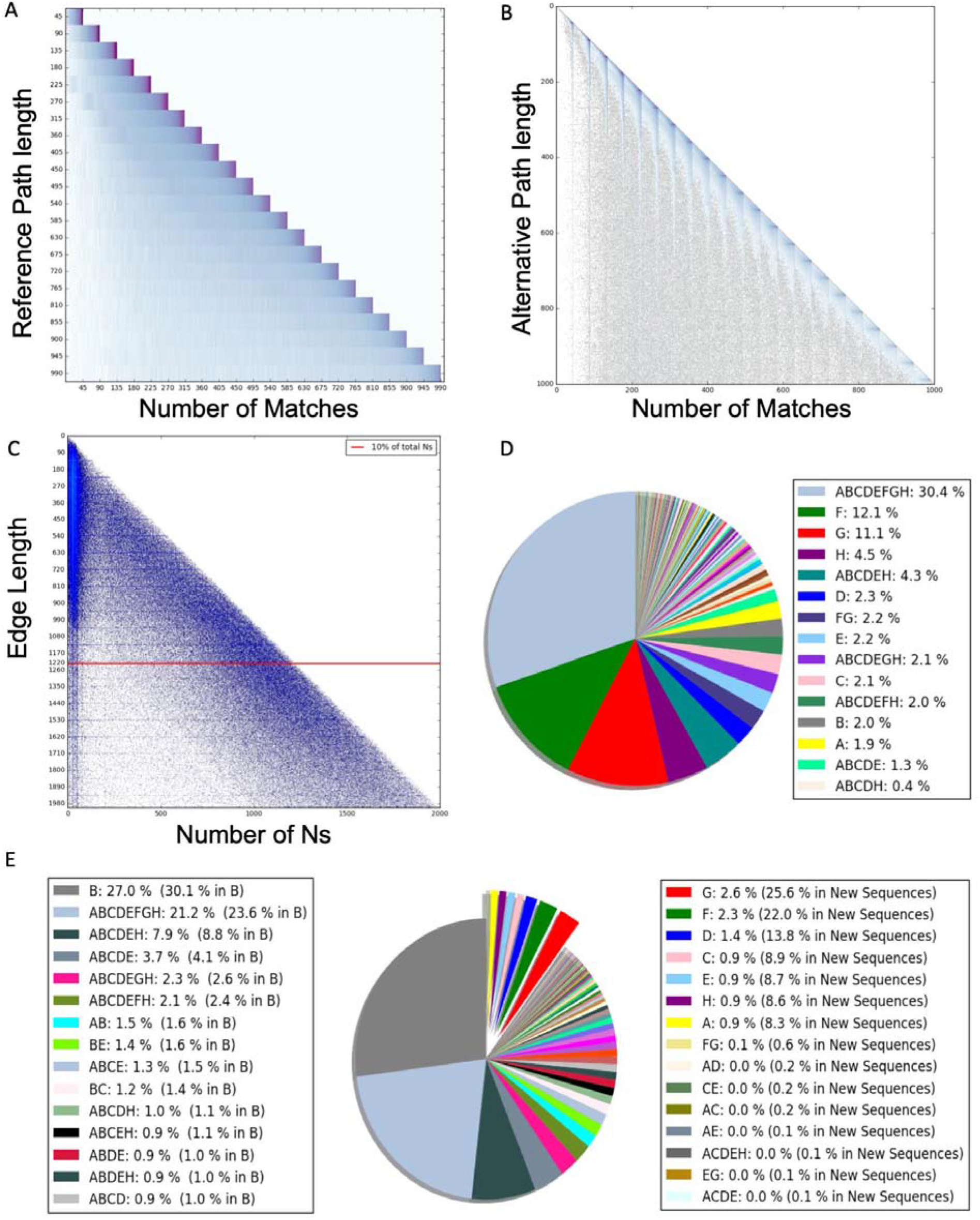
Sequence Sharing and Diversity. **A)** A two-dimensional histogram of alignment matches between alternate and reference parallel paths versus the total length of the reference path. Here we consider only reference paths less than 1000 bp. The quantization visible in reference lengths is due to each reference path being a multiple of 45bp. The density is centered on the diagonal, suggesting substantial sharing sequence between parallel reference and alternate paths. **B)** A two-dimensional histogram of alignment matches versus the alternative path length. Since the reference path lengths are multiples of 45bp, the number of matches exhibits artifacts. Nonetheless, the density is still centered along the diagonal further illustrating substantial sequence sharing between parallel paths. However, as the length of the alternative-paths increase, the variance in the number of matches also increases. **C)** A 2D histogram illustrating ambiguous base statistics. The density of ‘N’ counts versus edge length (for edges less than 2000 bps) is shown in this figure. 10% of ‘N’s appear in edges less than 1220 bps as indicated by the red line. **D)** A pie chart of edge Strain Diversity Patterns (SDPs) in CCGG. The ‘strain’ attribute indicates the founder genomes that share an edge, there are 255 distinct possibilities. We plot the fraction of edges that exhibit the same pattern of sharing. The top 15 SDPs comprise 80.9% of sharing patterns and their fractions are given in the legend. **E)** Illustrates the percentage of new base pairs in the CCGG according to SDP. The number of new base pairs that differ from the reference genome excluding Ns are plotted. By new bases, we refer to mismatches (SNPs) and insertions relative to the reference based on alignments of parallel edges. For long edges (> 1000bp), we approximate the number of new bases by the difference in lengths between alternate and reference paths. The pie chart includes 2,844,298,072 bp in total, 297,974,379 bp are new bases, which is 10.48% of the total size. This is an underestimate of the actual number of new bases due to our long edge approximation.

While anchor pairs separated by long gaps (more than 1000bp) in the standard mouse reference assembly comprise less than 5% of anchor intervals, they generally imply more sequence complexity. Possible reasons for these long gaps include genome assembly problems and large structural variants. For example, unresolved bases (*N*s) are overrepresented in long edges. A 2Dhistogram of the edge length (less than 2000 bps) versus the number of Ns in the edge’s sequence is shown in Figure 3C. Of the total 2,712,033,616 Ns in the CCGG, 742,627,244 Ns (27.4%) fall along these shorter edges. Edges less than 1224 bps (shown as a red line) explain 10% of Ns, with the remaining 90% of Ns lying on longer edges.

Long gaps between anchors often imply large-scale structural variants, including large deletions (in either a founder or CC sample), large insertions, or duplications that differentiate strains in these regions. Morgan *et al* reported 1749 *copy-number variants(CNVs)* and their coordinates in the CC founder strains as determined by sequencing of Diversity Outbred (DO) samples (Morgan et al. 2017). The DO population is related is derived from the CC and shares the same eight founders [ref]. We mapped these *CNVs* onto the CCGG and found that 71.58% (1252 in 1749) fall within a single long anchor gap. The average gap length (reference path length between adjacent anchor pairs) of the reported CNVs is 752,856 bp with 88.16% of them are longer than 1000 bp, while more than 90% of the total gaps are less than 1000 bp in the whole genome (Figure 2B). Thus, the reported CNVs are overrepresented in the long gap regions. We found that the alternate path lengths in long gaps varies more than in short gaps. We used path length from anchor to anchor as a measure rather than edge lengths, since floating nodes are common within long gaps. As shown in Figure 2, parts D and E, the density of reference path lengths versus alternative path lengths is more dispersed for long gaps as compared short gaps. Figure 2E also illustrates a cluster of large deletions on the alternative paths, where the reference path more than 6 Kb larger than alternative paths. This suggests that long gaps frequently represent sizable insertions or deletions. Thus, the CCGG points represents large scale variable regions for further biological exploration as well as areas where existing assemblies might require revisions.

The sharing patterns of CCGG edges among strains also provide interesting insights. We analyzed Strain Diversity Patterns (SDPs) of sequences based on founder strain annotations in the CCGG. Since anchors represent sequences shared by all 8 founder genomes, we treat them equivalently to an edge shared by all eight founders. The observed proportions of the 255 possible SDP patterns are shown in Figure 3B. The Top 15 SDPs and their fractions are illustrated in the legend. Edge sequences shared by all 8 founders are the most common in the CCGG and comprise 30.3% of the sequences; Private SDP patterns for the wild-derived mouse strains, CAST and PWK, contribute 12.1% and 11.1% of CCGG edges respectively; sequences that shared by all classical laboratory mouse strains contribute 4.3% of CCGG edges. These results correspond to the phylogeny of the 8 founders(Yang et al. 2011; Collaborative Cross 2012; Morgan and Welsh 2015). We next used the CCGG to estimate the number of new bases provided in the CCGG pangenome that differ from the standard mouse reference (GRC38). By new bases, we refer to mismatches and insertions that are not present in the reference assembly excluding Ns. For long gaps we make a conservative approximation of the number of additional bases as the difference in path lengths between alternative and the reference paths, excluding any Ns, since alignments in these regions are typically ambiguous. As shown in Figure 3E, we compared the SDP patterns in terms of reference-genome bases and the new bases. The pie chart includes 2,844,214,470 bp in total, 297,890,777 (10.47%) of which are new bases. As expected, private founder sequences contribute more than 95.9% of these new bases, and the wild-derived strains (PWK and CAST) alone contribute 47.6%. Of the SDP patterns that include the reference, the most common pattern are those private to C57BL/6J (B, 27.0% of CCGG bases, 30.1% of GRC38 bases), followed by conserved patterns shared by all 8 founder genomes (ABCDEFGH, 21.2% of CCGG bases, 23.6% of GRC38 bases), and then SDP patterns shared between *Mus musculus domesticus* haplotypes (ABCDEH, 7.9% of CCGG bases, 8.8% of GRC38 bases), followed by sequences common to all common laboratory mouse strains (ABCDE, 3.7% of CCGG bases, 4.1% of GRC38 bases). Overall, these CCGG statistics provide and overview of the actual sequence sharing and genetic diversity of the CC population.

### Investigating Genomic Features in the Graphical Genome

The CCGG represents genome features with overlaid annotations on both shared and private edges. We provide gene, exon, and interspersed repeat element annotations from repeat masker (Smit et al. 2015) on the mouse reference assembly path (B), and annotate their intervals using CIGAR strings (Fig S3). On average, a gene spans 110 anchor nodes, and an exon interval covers 1.58 anchor nodes. Since anchor sequences are conserved, we expected them to be enriched in genes. We found that 51.57% (4,751,256 of 9,212,137) of anchors fall within or overlap genes (compared to 42.60% of the Mus reference genome length that falls within one or more genes) and 4.55% of anchors (419,369 in 9,212,137) fall into or overlap exons (compared to 2.64% of the reference genome, which is overlaps exons).

The CCGG provides an effective resource for multi-genome comparisons by identifying strain-specific variants, especially in regions where variants that are difficult to characterize in a standard linear reference assembly. The graph structure places variants in the context of their haplotype, facilitating visualization of overall genome structures and comparisons of features. The CCGG provides an efficient means for identifying variants by traversing the graph and constructing the sub-paths between those annotated regions attributed to a certain *strain*. The number of paths between anchor pairs indicate the number of distinct haplotypes within the given region. If only a single path passes through the annotated region, it is conserved within the population. For the regions with multiple paths, one can further compare the sub-path lengths to determine if any indels fall in the regions. Most of the SNPs and tandem indels can be located by scanning through the *variant* cigar string on each edge.

The overall structure of gene *Dusp18* is shown in Figure 4A as an example illustrating conserved and variable genomic regions. A conserved region, which is indicated by the single path between anchor nodes A11.00086609 and A11.00086614 (3,897,406bp - 3,897,631bp in GRC38), encompasses the exon ENSMUSE00000682313. A highly variable region of the gene lies between CCGG anchors A11.00086572 and A11.00086576 (3,895,741bp - 3,895,921bp) has 8 distinct haplotypes. We examined all TSL1 annotated transcripts (Kinsella et al. 2011) of genes to estimate the number of distinct founder sub-paths within each. We found 95.94% of genes have multiple paths, suggesting that at least one variant exists in most genes. Moreover, 53.14% of genes have 8 paths, indicating a distinct haplotype for each founder. In coding region, we found that 44.3 % exons have a single common sequence, suggesting exons tend to be more conserved. However, the 55.69% of exons with multiple haplotypes suggests a substantial number of variants exist within coding regions.

**Figure 4.**
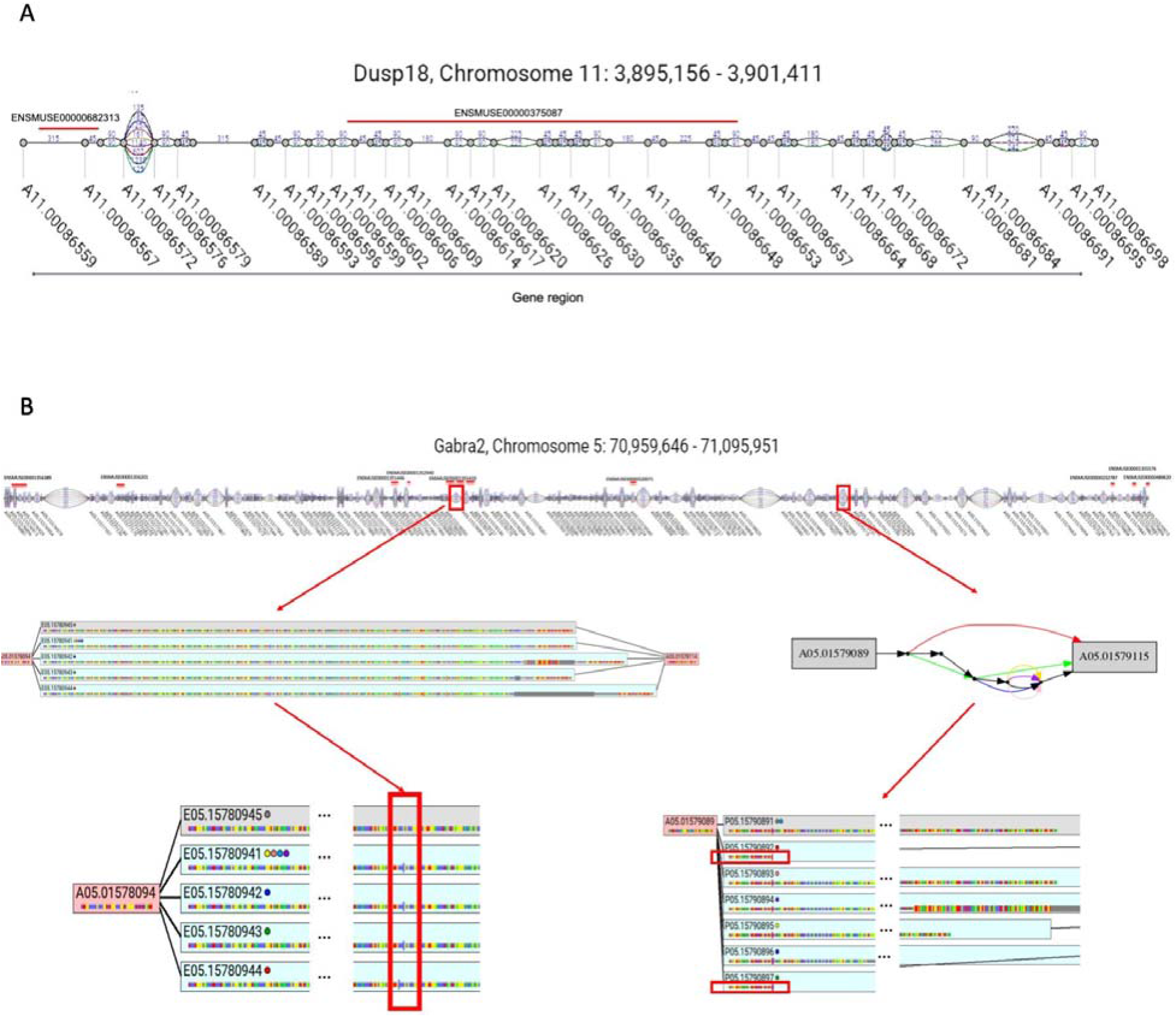
Gene Visualization and Variants Representation in CCGG. **A)** The genomic structure of *Dusp18* is shown. Highly variable regions and conserved regions can be identified in the graph, such as the interval between anchors A11.00086572 and A11.00086576 (3,895,741 - 3,895,921bp) with 8 haplotypes, as well as the conserved coding regions between A11.00086609 A11.00086614 (3,897,406bp - 3,897,631bp) in exon ENSMUSE00000682312. **B) A C57BL/6J specific functional variant** in Gabra2. The B6J specific functional deletion exists between anchors A05.01578094 and A05.01578114. In this gap, the reference path (B, grey box) is private, indicating a B6J specific feature in this region. By further examining the “variants” annotation for each path, a B6J specific deletion can be identified: all other paralleled path have the insertion of “A” in the same region. Another structural deletion in PWK and CAST within the Gabra2 gene body (71,059,388-71,059,864 bp) was reported by Sanger Institute. According to the reported region, we located this deletion between anchor A05.01579089 and A05.01579115 in CCGG. The graph shows that both the PWK and CAST path contain private features and their lengths are significantly shorter than the reference path, which is consistent with the previous report. Overall, CCGG provides an intuitive means for genome structure visualization and comparison.

Figure 4B illustrates variants in the *Gabra2* gene body. A recent study reported a C67BL/6J specific, functional non-coding variants in the *Gabra2* gene body (Mulligan et al. 2019). The mutation is a single base-pair deletion that became fixed in C67BL/6J between 1976 and 1991. We examined the CCGG subgraph containing *Gabra2* (Figure 4B) and found 387 anchors in gene’s region and 27 anchors fall within exons. Given the position reported by Mulligan et al, we located the variant to the interval between anchors A05.01578094 and A05.01578114. We found that there are 5 distinct paths between these two anchors and the reference path (B path) is private, suggesting a C57BL/6J specific sequence in this region. In addition, located about 411 base pairs distal of anchor A05.01578094, all the parallel edges share an insertion of “A”, indicating a deletion of “A” in the reference. We further identified indels exist in 141 anchor intervals within the gene, among these 10 intervals fall in coding regions. Differences in lengths between parallel and reference path length indicate potential insertions or deletions. A deletion located in the CCGG between anchors A05.01579089 and A05.01579115 where 7 distinct haplotypes exist in this homologous region and the path length of PWK and CAST haplotypes are significantly shorter than other genomes (Figure 6B). This is consistent with the Sanger Institute report that a structure variant (71,059,388-71,059,864 bp) exist within the *Gabra2* gene body in the wild-derived strains, PWK and CAST. Furthermore, the CCGG provides additional information. For example, in this same homologous region between anchors A05.01579089 and A05.01579115, the WSB haplotype is longer than the reference genome suggesting an insertion in WSB.

### CC Recombinations occur in more than 2000 gene regions

The Collaborative Cross is a multiparent recombinant inbred strain mouse panel derived from eight founder inbred strains. Each CC genome can be regarded as a mosaic of their 8 founder genomes and the haplotypes were reconstructed by imputation from the genome sequence of the corresponding founder inbred strain. Informative SNP were used to estimate the probability that a given genome region descended from each of the 8 founder strains (Srivastava et al. 2017). We annotated the CC paths in the graphical genome based on the founder probabilities determined from the genome sequencing data (see Method). By traversing the inferred founder paths of a CC sample from Srivastava et. al in the CCGG (in both forward and backward directions), we determined the distribution of recombination events per 2Mb interval (Figure 5A). We found 7353 total recombination over the entire genome. On average, recombination occur every 358190 bp (roughly 3 per Mb). We found that recombination events were enriched at the end of each chromosome and verified previously reported recombination hotspots and coldspots (Liu et al. 2014; Morgan et al. 2017).

**Figure 5.**
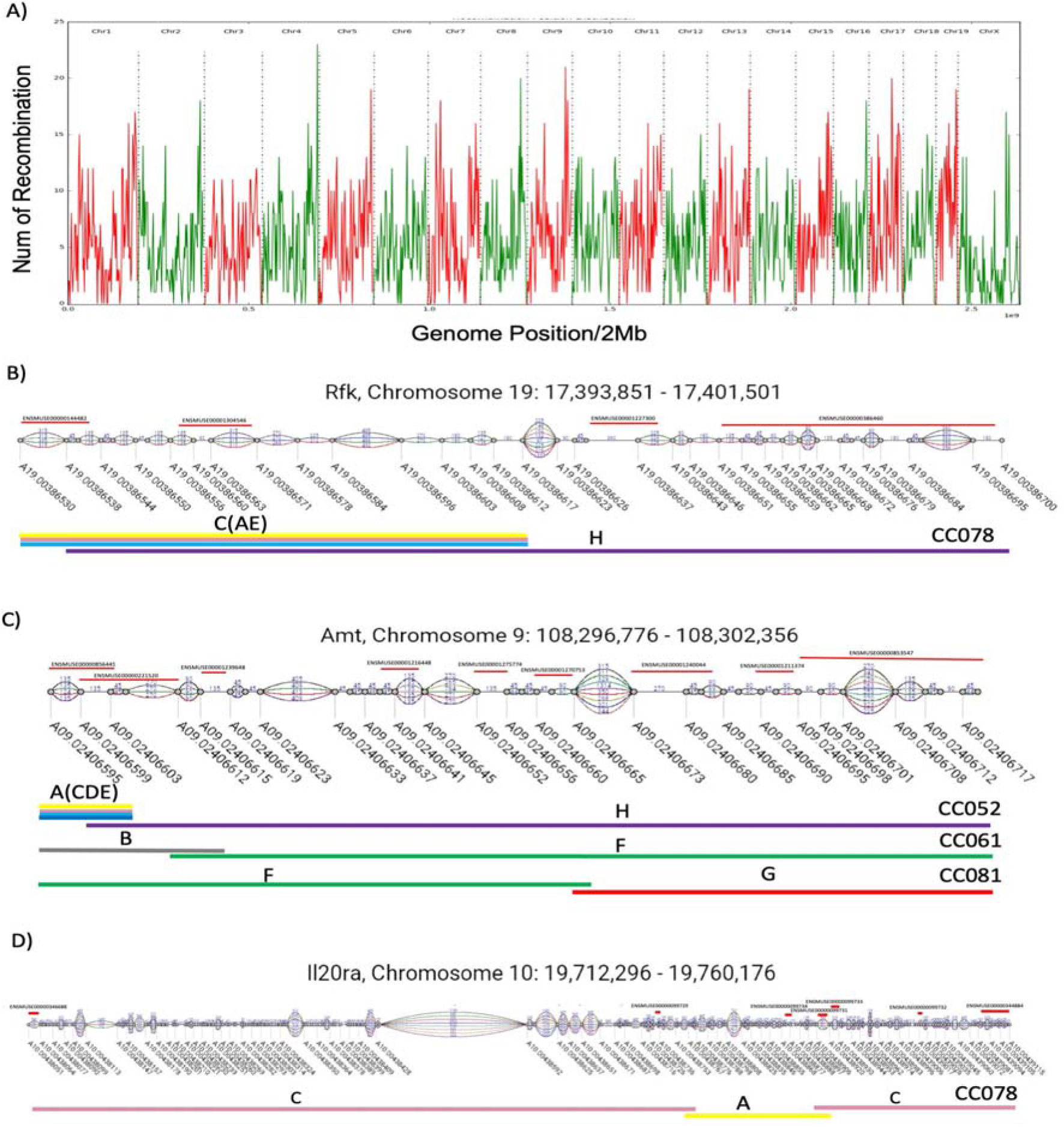
CC Recombination in genes and exons. **A)** The number of recombination in CC samples is plotted per 2Mb region. Recombination events are enriched at the ends of each chromosome. Recombination hotspots are also apparent. **B)** An example of recombination within a gene body. CC078 strain has a recombination within *Rfk* on chromosome 19. Exons lie on both sides of the recombination boundary which could generate novel isoforms in the CC strain. **C)** Multiple CC strains have recombination within a single gene. We found out CC052, CC061 and CC081 have recombination within the *Amt* gene. **D)** CC078 has two recombination events (C-A-C) within a single gene *Il20ra* that impacts a single exon. A total of 50 multiple recombination occur in 44 different genes. These mosaic genes might lead to functional consequences.

We further analyzed the recombination events within annotated functional regions, such as gene bodies. We considered 46036 TSL1 gene regions in all 75 CC strains. We found out 3293 distinct recombination events occurred in the 2108 annotated gene regions in 75 CC strains, 50 of which contain multiple recombination events within a single gene; 96 distinct recombination events occurred in the 90 exon regions involving 82 genes in 50 CC strains. The recombination information within gene and exons is provided in Supplementary Table 5 and 6. As shown in Figure 5, CC078 has a recombination within *Rfk* gene body around chr19: 17M. We found that CC078 is of type C,129S1/SvImJ (which is indistinguishable from types A, A/J and E, NZO/HlLtJ) at the beginning of this gene, and transitions into haplotype H (WSB/EiJ) by the gene’s end. By traversing the graph both directions, we discovered that this transition occurs between anchor A19.00386617 and A19.00386538 (17,397,766 - 17,394,211 bp). We find exons exist both before and after this recombination and a WSB private path exists in an exon (ENSMUSE00000386460), which suggest that the recombination may lead to novel transcriptional isoforms for gene *Rfk* in CC078. Besides the 96 recombination events that occur within the exon regions, we found 2676 recombination events between exons from different founder strains. These new gene versions may affect the RNA splicing and produce novel isoforms.

Furthermore, recombination hotspots are identified in gene bodies. We find that 639 genes have more than one CC strain recombination within the same gene. As shown in Figure 5C, CC052, CC061 and CC081 have recombination within *Amt*. Exons exist both before and after the recombination boundaries in these strains, which might lead to gene functional diversity in those CC strains. Besides, five genes, *Fgfr2*, *Gm20388*, *Osbpl10*, *Pde4d*, and *Prkg1* contain more than 10 recombination events in different CC strains. Among them, *Gm20388* (4434881 bp) contains the largest number of recombination involving 17 distinct CC strains. These results suggest genes that potentially might express new isoforms in CC population and are interesting targets for further biological exploration.

The recombination within gene bodies can be complex. Several CC strains have multiple recombination events within a single gene body. For example, CC078 transitions from haplotype C (129S1/SvImJ) to A (A/J) and back to C (129S1/SvImJ) within gene *Il20ra* (Figure 5D). We found 50 multi-recombination events within a single gene, occurring in 44 distinct genes. Among them, 13 strains have recombination events impacting more than one gene body, suggesting the genomes of these strains could have more complex gene structure. For example, CC058 has multiple recombination in 4 different genes, including *Galnt13*, *Kif21a*, *Gm10556*, and *Slit1*. Multiple recombination within the same gene could lead to novel isoforms and other functional consequences.

Overall, the CCGG provides a novel resource combining genomic data from multiple linear genome assemblies. It enables the exploration of genomic variants and recombination events across different samples. This is of potential interest to be generalized in multi-genomic analysis and comparative study for genomic structure analysis.

## Discussion

In this paper, we introduce a pangenome resource for searching, annotating, and comparing genomic assemblies from a widely used mouse genetic reference population, collaborative cross. It combines 8 founder and 75 CC genomes with 2091 contigs into a single graph representation, which directly captures important genetics aspects such as identity-by-descent, and highly variable genomic regions. Our graph representation introduces a special class of conserved nodes, which we call *anchors* that organizes genomes into disjoint homologous segments and provides a new framework for labeling genomic regions while maintaining *monotonicity, backward-compatibility,* and *uniformity.* It also allows annotations to be placed in multiple genomic contexts and supports incremental updates as the assembly is improved without needing to re-establish all other annotations. The genome of each mouse strain represented in the CCGG can be extracted as classical linear genome and used in the standard bioinformatics pipelines. The graph structure also provides an effective way for searching, comparing and visualizing genomic features and facilitates the tool chain development for downstream analysis. The properties of graphical genome, such as the haplotype sequence diversity and gap lengths are informative for further biological discovery, including the identification of structural variants and correlation analysis. Combining the inferred CC haplotype with graphical genome facilitates understanding of recombination events and enables the functional analysis of genes beyond point mutations. Lastly, the CCGG is a versatile representation of genomic sequences, and provides a significant space savings.

The CCGG can readily be extended to incorporate new inbred strains and other recombinant inbred lines. For example, by including a single strain, DBA/2J, the utility of the CCGG can be expanded to include nearly 100 additional BXD strains (Taylor et al. 1999; Peirce et al. 2004; Philip et al. 2010). The CCGG is also useful for characterizing the Diversity Outcross (DO) population (Svenson et al. 2012), an outbred population derived from the same set of CC founders developed for their fine-mapping potential.

Overall, CCGG presents a holistic view of sequence diversity in mouse species and provides an alternative genome model for the development of future bioinformatic tools and biological assays. The genomic sequence diversity of the CCGG clearly illustrates that relying on a single linear reference genome derived from a single individual is not sufficient for alignment and reference-specific variants detection, and that multi-strain sequencing need to be applied to reference pangenomes construction to understand the genetic basis of a population. The CCGG proposed in this work contributes for Mus reference pangenome development. It served as a substrate for further sequence content improvement, CC private sequence assembly, and downstream analysis tool-chain development.

### Potential new tool chains based on CCGG

We suggest that it is both possible and informative to perform short-read sequence alignments directly onto segments of the CCGG. This could be accomplished efficiently by constructing an msBWT of anchor-edge-anchor extended into flanking edges by some constant amount depending on the maximum allowed read-length. This would allow every read fragment to be mapped to one or more gaps between anchors. The resulting msBWT and FM-index for the CCGG would likely be only slightly larger than a BWT index constructed from a traditional reference assembly. It also is possible to add special edge types into the CCGG representing annotated splice junctions for use in RNA alignments.

The CCGG can also be used for genomic comparisons between CC strains and to further refine the recombination boundaries. The chain of node-edge-node’s along a path provides a high-level abstraction of the genome that allows synteny to compare between paths much more efficiently than direct sequence alignments. These descriptions can be compared directly between strains to assess their similarities and differences at a high level. Combining with comparative phylogenetic methods, much informative illustrations for the CC genealogy can be obtained from the graphical genome with multi-sequence assemblies.

We also propose that association mapping methods can be developed using the CCGG to find regions where patterns of genomic variation correlate to phenotypes. Collaborative Cross serves a community resource for genetic analysis of complex traits. Most of genome wide association studies focus on SNPs due to the difficulty of recording and identifying structural variants. The CC graphical genome provides an effective way to identify potential structural variant regions and provide a platform for developing association study at whole genome level.

### Improving the sequence content of the CCGG

The sequence content of the draft version of CCGG comes from the 8 founder reference assemblies. However, we expect that CC lines will introduce their private edges in the graph due to *de novo* mutations that have fixed within a strain. Examples include the deletions reported in (Srivastava et al. 2017). Moreover, there are many unresolved genomic regions, represented by runs of N’s, in the 7 new assemblies from Lilue et al. 2018, where the CC sequence data provides significantly more power to resolve. On average, there are more than eight CC strains representing each of the eight founders at every place in the genome. This is equivalent to more than 160x genomic coverage, from biological replicates, which should greatly reduce the impact of private variants from any one sample. Furthermore, the genomic assembly process is greatly simplified by the inclusion of anchor sequences that serve as seeds for local assembly tasks.

On the other hand, much of the X and Y chromosome sequences are repetitive (Mueller et al. 2008). The natural implication from this fact is that there will be less anchor nodes in the X and Y chromosome. For chromosome X (171031299 bp), the number of anchors is 89.5% less than the comparably sized chromosome 2 (182113224 bp) (Table 1). As shown in Figure. 2B, the cumulative distribution of anchor distances on X is significantly different from the distribution in autosomes. These results suggest that chromosome X has relatively longer edges than autosomes. The released versions of the new genome assemblies from (Lilue et al. 2018) do not even include sequence for Y. Thus, we constructed a draft chromosome Y graph using unique 45-mers present in all male CC samples. We constructed a single edge separating each pair of anchors. These edges are currently shared on all paths and are populated with sequence from the mouse reference assembly. It stands as a draft for future work on assembling better models for Y in CC strain genomes.

## Methods

This section highlights the key components in the CCGG construction and the primary sources of data. More details about the construction method are provided in the supplementary material.

### Sequence content and 45-mer occurrence matrices

The sequence contents of CCGG were constructed by merging the standard mouse reference (Mouse Genome Sequencing et al. 2002; Church et al. 2009) with *de novo* assemblies of the 7 inbred mouse genomes assembled by Lilue et al. 2018 (Lilue et al. 2018). The first stage of the CCGG’s construction focused on identifying a set of conserved and unique sequences to establish a set of genomic anchors. This began by constructing Multi-string Burrows-Wheeler Transforms (msBWTs) for all lanes and pair ends of the Illumina read sets from (Srivastava et al. 2017) (Srivastava et al. 2017) and (Keane et al. 2011) using the package from (Holt et al. 2013). Next, an occurrence-count matrix for every non-overlapping 45-mers from the standard mouse reference in each of the 69 sequenced CC samples reported in Srivastava et al. 2017 and 8 additional CC samples from Shorter et al. 2019.

### Annotation

The annotations in CCGG include genes, exons, repeat masks, variants and CC hyplotytes. The start and end coordinates of gene and exon intervals of TSL1 transcripts in the mouse reference genome were obtained from Ensembl Biomart (Kinsella et al. 2011). The repeat-masker intervals for the standard mouse reference were obtained from http://www.repeatmasker.org/. The variants annotation was obtained by aligning the parallel alternative sequences to the reference path using the same end-to-end scoring function in Bowtie2 (Langmead and Salzberg 2012). The CC haplotypes were previously inferred from CC sequencing data using forward-backward hidden Markov model (Srivastava et al.2017) and the annotation was added by viewing CC genomes as a mosaic of their 8 founder sequences. Overall, 46063 genes, 726152 exons, 5700130 repeatmasker regions and variants were annotated and approximate genomes for 75 CC strains based on their haplotype annotations were provided.

### Compression

CCGG was compressed by adding floating nodes within gaps where any path between anchor pairs exceeded 1000bp. We partitioned the sequences in those long gaps into non-overlapping 45-mers (in the case where sequence lengths are not multiples of 45 bp, we allowed for overlaps in the middle). Each 45-mer is assigned with a unique index based on its sequence content. Based on these indices, we were able to find shared 45-mers among parallel edges. We merged consecutive shared 45-mers and inserted floating nodes at the beginning and the end of the shared sequences. The floating nodes themselves don’t carry any sequence content or annotation, so the compression was achieved by merging sequences between floating nodes.

## Data Access

The Collaborative Cross Graphical Genome source code and data is available at https://github.com/jay1723/GraphicalGenome. It includes the CCGG as a collection of FASTA files representing the graphical pangenome, a Python-based API for loading, navigating, annotating, and saving the pangenome, and a command-line tool for extracting paths (traditional linear genome assemblies) from the CCGG.

The CCGG sequence files can be downloaded from https://devel.csbio.unc.edu/graphicalGenome. For convenience, separate FASTA files are provided for each chromosome’s set of anchor nodes and edges.

## Acknowledgements

The motivation for constructing the CCGG is a direct result of requests for better genomic models from many collaborators using the Collaborative Cross, including Darla Miller, Gary Churchill, Martin Ferris, Samir Kelada, Lisa Tarantino, William Valdar, David Aylor, and David Threadgill. This work is supported by a resource development grant from NIH/NHGRI U24-HG010100, to develop genomic resources for the Collaborative Cross.

## Disclosure Declaration

All authors declare no conflict of interests.

## Notes

http://devel.csbio.unc.edu/GraphicalGenome/index.html

## References

Aylor DL, Valdar W, Foulds-Mathes W, Buus RJ, Verdugo RA, Baric RS, Ferris MT, Frelinger JA, Heise M, Frieman MB et al. 2011. Genetic analysis of complex traits in the emerging Collaborative Cross. Genome Res 21: 1213–1222.

Baier U, Beller T, Ohlebusch E. 2015. Graphical pan-genome analysis with compressed suffix trees and the Burrows–Wheeler transform. Bioinformatics 32: 497–504.

Bottomly D, Ferris MT, Aicher LD, Rosenzweig E, Whitmore A, Aylor DL, Haagmans BL, Gralinski LE, Bradel-Tretheway BG, Bryan JT et al. 2012. Expression quantitative trait Loci for extreme host response to influenza a in pre-collaborative cross mice. G3 (Bethesda) 2: 213–221.

Church DM, Goodstadt L, Hillier LW, Zody MC, Goldstein S, She X, Bult CJ, Agarwala R, Cherry JL, DiCuccio M et al. 2009. Lineage-specific biology revealed by a finished genome assembly of the mouse. PLoS Biol 7: e1000112.

Church DM, Schneider VA, Graves T, Auger K, Cunningham F, Bouk N, Chen HC, Agarwala R, McLaren WM, Ritchie GR et al. 2011. Modernizing reference genome assemblies. PLoS Biol 9: e1001091.

Church DM, Schneider VA, Steinberg KM, Schatz MC, Quinlan AR, Chin C-S, Kitts PA, Aken B, Marth GT, Hoffman MM. 2015. Extending reference assembly models. Genome biology 16: 13.

Collaborative Cross C. 2012. The genome architecture of the Collaborative Cross mouse genetic reference population. Genetics 190: 389–401.

Computational Pan-Genomics C. 2018. Computational pan-genomics: status, promises and challenges. Brief Bioinform 19: 118–135.

Dilthey A, Cox C, Iqbal Z, Nelson MR, McVean G. 2015. Improved genome inference in the MHC using a population reference graph. Nature genetics 47: 682.

Gartner F, Honer Zu Siederdissen C, Muller L, Stadler PF. 2018. Coordinate systems for supergenomes. Algorithms Mol Biol 13: 15.

Herbig A, Jager G, Battke F, Nieselt K. 2012. GenomeRing: alignment visualization based on SuperGenome coordinates. Bioinformatics 28: i7–15.

Hogg JS, Hu FZ, Janto B, Boissy R, Hayes J, Keefe R, Post JC, Ehrlich GD. 2007. Characterization and modeling of the Haemophilus influenzae core and supragenomes based on the complete genomic sequences of Rd and 12 clinical nontypeable strains. Genome Biol 8: R103.

Holt J, Huang S, McMillan L, Wang W. 2013. Read annotation pipeline for high-throughput sequencing data. In Proceedings of the International Conference on Bioinformatics, Computational Biology and Biomedical Informatics, p. 605. ACM.

Hu Z, Sun C, Lu KC, Chu X, Zhao Y, Lu J, Shi J, Wei C. 2017. EUPAN enables pan-genome studies of a large number of eukaryotic genomes. Bioinformatics 33: 2408–2409.

Huang S, Holt J, Kao C-Y, McMillan L, Wang W. 2014. A novel multi-alignment pipeline for high-throughput sequencing data. Database 2014.

Jacobsen A, Hendriksen RS, Aaresturp FM, Ussery DW, Friis C. 2011. The Salmonella enterica pan-genome. Microb Ecol 62: 487–504.

Keane TM, Goodstadt L, Danecek P, White MA, Wong K, Yalcin B, Heger A, Agam A, Slater G, Goodson M et al. 2011. Mouse genomic variation and its effect on phenotypes and gene regulation. Nature 477: 289–294.

Kelada SN, Aylor DL, Peck BC, Ryan JF, Tavarez U, Buus RJ, Miller DR, Chesler EJ, Threadgill DW, Churchill GA et al. 2012. Genetic analysis of hematological parameters in incipient lines of the collaborative cross. G3 (Bethesda) 2: 157–165.

Kinsella RJ, Kähäri A, Haider S, Zamora J, Proctor G, Spudich G, Almeida-King J, Staines D, Derwent P, Kerhornou A. 2011. Ensembl BioMarts: a hub for data retrieval across taxonomic space. Database 2011.

Langmead B, Salzberg SL. 2012. Fast gapped-read alignment with Bowtie 2. Nat Methods 9: 357–359.

Lapierre P, Gogarten JP. 2009. Estimating the size of the bacterial pan-genome. Trends Genet 25: 107–110.

Lefebure T, Stanhope MJ. 2007. Evolution of the core and pan-genome of Streptococcus: positive selection, recombination, and genome composition. Genome Biol 8: R71.

Leist SR, Pilzner C, van den Brand JM, Dengler L, Geffers R, Kuiken T, Balling R, Kollmus H, Schughart K. 2016. Influenza H3N2 infection of the collaborative cross founder strains reveals highly divergent host responses and identifies a unique phenotype in CAST/EiJ mice. BMC Genomics 17: 143.

Li R, Li Y, Zheng H, Luo R, Zhu H, Li Q, Qian W, Ren Y, Tian G, Li J et al. 2010. Building the sequence map of the human pan-genome. Nat Biotechnol 28: 57–63.

Lilue J, Doran AG, Fiddes IT, Abrudan M, Armstrong J, Bennett R, Chow W, Collins J, Collins S, Czechanski A et al. 2018. Sixteen diverse laboratory mouse reference genomes define strain-specific haplotypes and novel functional loci. Nat Genet 50: 1574–1583.

Liu EY, Morgan AP, Chesler EJ, Wang W, Churchill GA, Pardo-Manuel de Villena F. 2014. High-resolution sex-specific linkage maps of the mouse reveal polarized distribution of crossovers in male germline. Genetics 197: 91–106.

Manet C, Simon-Loriere E, Jouvion G, Hardy D, Prot M, Conquet L, Flamand M, Panthier JJ, Sakuntabhai A, Montagutelli X. 2019. Genetic diversity of Collaborative Cross mice controls viral replication, clinical severity and brain pathology induced by Zika virus infection, independently of Oas1b. J Virol doi:10.1128/JVI.01034-19.

Marcus S, Lee H, Schatz MC. 2014. SplitMEM: a graphical algorithm for pan-genome analysis with suffix skips. Bioinformatics 30: 3476–3483.

McCarthy CGP, Fitzpatrick DA. 2019. Pan-genome analyses of model fungal species. Microb Genom 5.

Morgan AP, Gatti DM, Najarian ML, Keane TM, Galante RJ, Pack AI, Mott R, Churchill GA, de Villena FP-M. 2017. Structural variation shapes the landscape of recombination in mouse. Genetics 206: 603–619.

Morgan AP, Welsh CE. 2015. Informatics resources for the Collaborative Cross and related mouse populations. Mamm Genome 26: 521–539.

Mouse Genome Sequencing C Waterston RH Lindblad-Toh K Birney E Rogers J Abril JF Agarwal P Agarwala R Ainscough R Alexandersson M et al. 2002. Initial sequencing and comparative analysis of the mouse genome. Nature 420: 520–562.

Mulligan MK, Abreo T, Neuner SM, Parks C, Watkins CE, Houseal MT, Shapaker TM, Hook M, Tan H, Wang X et al. 2019. Identification of a Functional Non-coding Variant in the GABA A Receptor alpha2 Subunit of the C57BL/6J Mouse Reference Genome: Major Implications for Neuroscience Research. Front Genet 10: 188.

Munger SC, Raghupathy N, Choi K, Simons AK, Gatti DM, Hinerfeld DA, Svenson KL, Keller MP, Attie AD, Hibbs MA et al. 2014. RNA-Seq alignment to individualized genomes improves transcript abundance estimates in multiparent populations. Genetics 198: 5973.

Paten B, Novak AM, Eizenga JM, Garrison E. 2017. Genome graphs and the evolution of genome inference. Genome Res 27: 665–676.

Peirce JL, Lu L, Gu J, Silver LM, Williams RW. 2004. A new set of BXD recombinant inbred lines from advanced intercross populations in mice. BMC Genet 5: 7.

Peter J, De Chiara M, Friedrich A, Yue JX, Pflieger D, Bergstrom A, Sigwalt A, Barre B, Freel K, Llored A et al. 2018. Genome evolution across 1,011 Saccharomyces cerevisiae isolates. Nature 556: 339–344.

Philip V, Duvvuru S, Gomero B, Ansah T, Blaha C, Cook M, Hamre K, Lariviere W, Matthews D, Mittleman G. 2010. High-throughput behavioral phenotyping in the expanded panel of BXD recombinant inbred strains. Genes, Brain and Behavior 9: 129–159.

Philip VM, Sokoloff G, Ackert-Bicknell CL, Striz M, Branstetter L, Beckmann MA, Spence JS, Jackson BL, Galloway LD, Barker P et al. 2011. Genetic analysis in the Collaborative Cross breeding population. Genome Res 21: 1223–1238.

Rand KD, Grytten I, Nederbragt AJ, Storvik GO, Glad IK, Sandve GK. 2017. Coordinates and intervals in graph-based reference genomes. BMC Bioinformatics 18: 263.

Shorter JR, Najarian ML, Bell TA, Blanchard M, Ferris MT, Hock P, Kashfeen A, Kirchoff KE, Linnertz CL, Sigmon JS. 2019. Whole Genome Sequencing and Progress Toward Full Inbreeding of the Mouse Collaborative Cross Population. G3: Genes, Genomes, Genetics 9: 1303–1311.

Smit A, Hubley R, Green P. 2015. RepeatMasker Open-4.0. 2013–2015.

Srivastava A, Morgan AP, Najarian ML, Sarsani VK, Sigmon JS, Shorter JR, Kashfeen A, McMullan RC, Williams LH, Giusti-Rodriguez P et al. 2017. Genomes of the Mouse Collaborative Cross. Genetics 206: 537–556.

Stevenson KR, Coolon JD, Wittkopp PJ. 2013. Sources of bias in measures of allele-specific expression derived from RNA-seq data aligned to a single reference genome. BMC genomics 14: 536.

Svenson KL, Gatti DM, Valdar W, Welsh CE, Cheng R, Chesler EJ, Palmer AA, McMillan L, Churchill GA. 2012. High-resolution genetic mapping using the Mouse Diversity outbred population. Genetics 190: 437–447.

Taylor BA, Wnek C, Kotlus BS, Roemer N, MacTaggart T, Phillips SJ. 1999. Genotyping new BXD recombinant inbred mouse strains and comparison of BXD and consensus maps. Mammalian genome 10: 335–348.

Tettelin H, Masignani V, Cieslewicz MJ, Donati C, Medini D, Ward NL, Angiuoli SV, Crabtree J, Jones AL, Durkin AS et al. 2005. Genome analysis of multiple pathogenic isolates of Streptococcus agalactiae: implications for the microbial “pan-genome”. Proc Natl Acad Sci U S A 102: 13950–13955.

Tettelin H, Riley D, Cattuto C, Medini D. 2008. Comparative genomics: the bacterial pangenome. Curr Opin Microbiol 11: 472–477.

Tian X, Li R, Fu W, Li Y, Wang X, Li M, Du D, Tang Q, Cai Y, Long Y. 2018. Generating a sequence map of the pig pan-genome. bioRxiv: 459453.

Yang H, Wang JR, Didion JP, Buus RJ, Bell TA, Welsh CE, Bonhomme F, Yu AH, Nachman MW, Pialek J et al. 2011. Subspecific origin and haplotype diversity in the laboratory mouse. Nat Genet 43: 648–655.

Zheng S, Cherniack AD, Dewal N, Moffitt RA, Danilova L, Murray BA, Lerario AM, Else T, Knijnenburg TA, Ciriello G et al. 2016. Comprehensive Pan-Genomic Characterization of Adrenocortical Carcinoma. Cancer Cell 30: 363.

Zhou Z, Lundstrom I, Tran-Dien A, Duchene S, Alikhan NF, Sergeant MJ, Langridge G, Fotakis AK, Nair S, Stenoien HK et al. 2018. Pan-genome Analysis of Ancient and Modern Salmonella enterica Demonstrates Genomic Stability of the Invasive Para C Lineage for Millennia. Curr Biol 28: 2420–2428 e2410.

